# Mitochondrial pseudogenes suggest repeated inter-species hybridization among direct human ancestors

**DOI:** 10.1101/134502

**Authors:** Konstantin Popadin, Konstantin Gunbin, Leonid Peshkin, Sofia Annis, Zoe Fleischmann, Genya Kraytsberg, Natalya Markuzon, Rebecca R. Ackermann, Konstantin Khrapko

**Author notes:** Corresponding author: Konstantin Khrapko Professor Department of Biology 134 Mugar Bldg 360 Huntington Avenue Northeastern University Boston MA 02115.

## Abstract

The hypothesis that the evolution of humans involved hybridization between diverged species has been actively debated in recent years. We present novel evidence in support of this hypothesis: the analysis of nuclear pseudogenes of mtDNA (“NUMTs”). NUMTs are considered “mtDNA fossils”, as they preserve sequences of ancient mtDNA and thus carry unique information about ancestral populations. Our comparison of a NUMT sequence shared by humans, chimpanzees, and gorillas with their mtDNAs implies that, around the time of divergence between humans and chimpanzees, our evolutionary history involved the interbreeding of individuals whose mtDNA had diverged as much as ~4.5 Myr prior. This large divergence suggests a distant interspecies hybridization. Additionally, analysis of two other NUMTs suggests that such events occurred repeatedly. Our findings suggest a complex pattern of speciation in primate human ancestors and provide a potential explanation for the mosaic nature of fossil morphology found at the emergence of the hominin lineage.

## Introduction

Increasingly, the emergence and evolution of our species is being revealed as a period characterized by genetic exchange between divergent lineages. For example, we now have evidence of hybridization between Neanderthals and people expanding from Africa (Sankararaman et al., 2012), (Vernot et al., 2016), between Denisovans and humans (Meyer et al., 2012), between Neanderthals and Denisovans (Prufer et al., 2014), and between Denisovans and an unidentified hominin (Prufer et al., 2014). *Hominin* here denotes human lineage upon its separation from the chimpanzee. The term *Hominine*, in contrast, denotes the human, chimpanzee and gorilla clade, upon its separation from the orangutan lineage (Ref needed). Denisovan-like mtDNA has also been detected in earlier (ca. 400,000ya) hominins (Meyer et al. 2014), implying mtDNA introgression or hybridization. Evidence also exists for gene flow (ca.35Kya) between African populations that diverged 700,000 years ago (Hammer et al., 2011). These studies indicate that hybridization was prevalent during the period of emergence of *Homo sapiens*, and suggest that it may be the rule rather than the exception in hominin evolution. However, we have little information on the presence or prevalence of hybridization during earlier (pre-1Ma) periods in human evolution. A study (Patterson et al., 2006) concluded that the hominin lineage first significantly diverged from the chimpanzee lineage, but later hybridized back, before finally diverging again. This study prompted an intense debate (Barton, 2006) (Innan and Watanabe, 2006), (Wakeley, 2008), (Patterson et al., 2008), (Presgraves and Yi, 2009), (Yamamichi et al., 2012), but has remained the sole piece of evidence for such an early admixture event. Although none of the subsequent studies fully rejected or confirmed the hybridization scenario, most pointed to the lack of sufficient evidence to uphold it. The research presented here supports a similar early hybridization scenario using an entirely different approach. Moreover, our analyses suggest that distant interbreedings occurred repeatedly among our distant ancestors.

The evidence for interspecies hybridization presented here comes from special type of pseudogenes (“NUMTs”) that are fragments of mitochondrial DNA (mtDNA) integrated into the nuclear genome. There are hundreds of NUMT sequences in the human genome (Ramos et al.). NUMTs found in the present day human genome have been inserting into nuclear DNA since tens of millions years ago, and this process continues today (Srinivasainagendra 2017) We have recently reported evidence that insertion of NUMTs into nuclear genome might have accelerated during the emergence of the genus Homo (Gunbin 2016). NUMTs are considered “DNA fossils”, since they preserve ancient mtDNA sequences virtually unchanged due to significantly lower mutational rate in the nuclear versus mitochondrial genome (Zischler et al., 1995). NUMTs therefore offer an opportunity to peek into the distant past of populations (Bensasson et al., 2001), (Baldo et al., 2011), (Wang et al., 2015). Here, we demonstrate that a NUMT on chromosome 5 descends from a mitochondrial genome that had been highly divergent from our ancestors’ mtDNA at the time of becoming a pseudogene. This implies that this pseudogene should have been created in an individual from a (hominine) species that at the time of insertion was highly diverged from our direct ancestor. For this pseudogene to end up in our genome, this (now extinct) hominine should have hybridized with our direct ancestors. Moreover, our analysis of additional NUMTs with similar phylogenic history, implies that this scenario was not unique.

## Results

### A NUMT on chromosome 5 originated from a highly divergent mitochondrial genome

In an early screen of human pseudogenes of mtDNA (Li-Sucholeiki et al., 1999) we discovered a pseudogene sequence on chromosome 5 which later turned out to be a large (~9kb) NUMT called here “ps5”. We then discovered close homologs of ps5 in the chimpanzee and gorilla genomes, i.e. in all contemporary *hominines*, but absent in orangutan and more distant primates (see **Suppl. Note 1**). This NUMT turned out to have an extraordinary evolutionary history.

A joint phylogenetic tree of the 3 ps5 NUMTs and the mtDNA sequences of great apes (**Fig. 1A**) has a very surprising shape. One would expect that, as selectively neutral loci, pseudogenes should approximately follow the evolutionary paths of the species in which they reside. That is, the NUMT sub-tree should resemble the mtDNA sub-tree, which is a good representation of the great ape evolution. One may expect, though, that all branches of the NUMT tree should be shorter than those of mtDNA tree, as mutation rate in the nuclear DNA is expected to be lower than in mtDNA. Contrary to these expectations, the NUMT has a very different shape: a very long stem (“ps5 stem”) and short branches. Even more intriguingly, phylogenetic mutational analysis (**Suppl. Note 6**) showed that the mutations of the ps5 stem contain a very high proportion of synonymous changes, similar to mtDNA branches. In contrast, mutations in the outer pseudogene branches (**Fig. 1A**, colored blue) contain a significantly higher proportion of non-synonymous changes (p<0.00005,), as expected for a truly pseudogenic, dysfunctional sequence. Thus ps5 sequence has been evolving under mitochondrial selective constraints, i.e. as a part of a functional mitochondrial genome, until it gave rise to a pseudogene which then split into the *Homo*, *Pan*, and *Gorilla* variants. The impressive length of the ps5 stem implies that at the time of it’s insertion in to the nuclear genome, the mtDNA predecessor of ps5 NUMT was highly divergent from the *Homo/Pan/Gorilla* ancestral mtDNA. Because the rate of evolution of mtDNA is relatively stable and well-documented, this divergence can be evaluated quantitatively with reasonable confidence.

**Figure 1.**
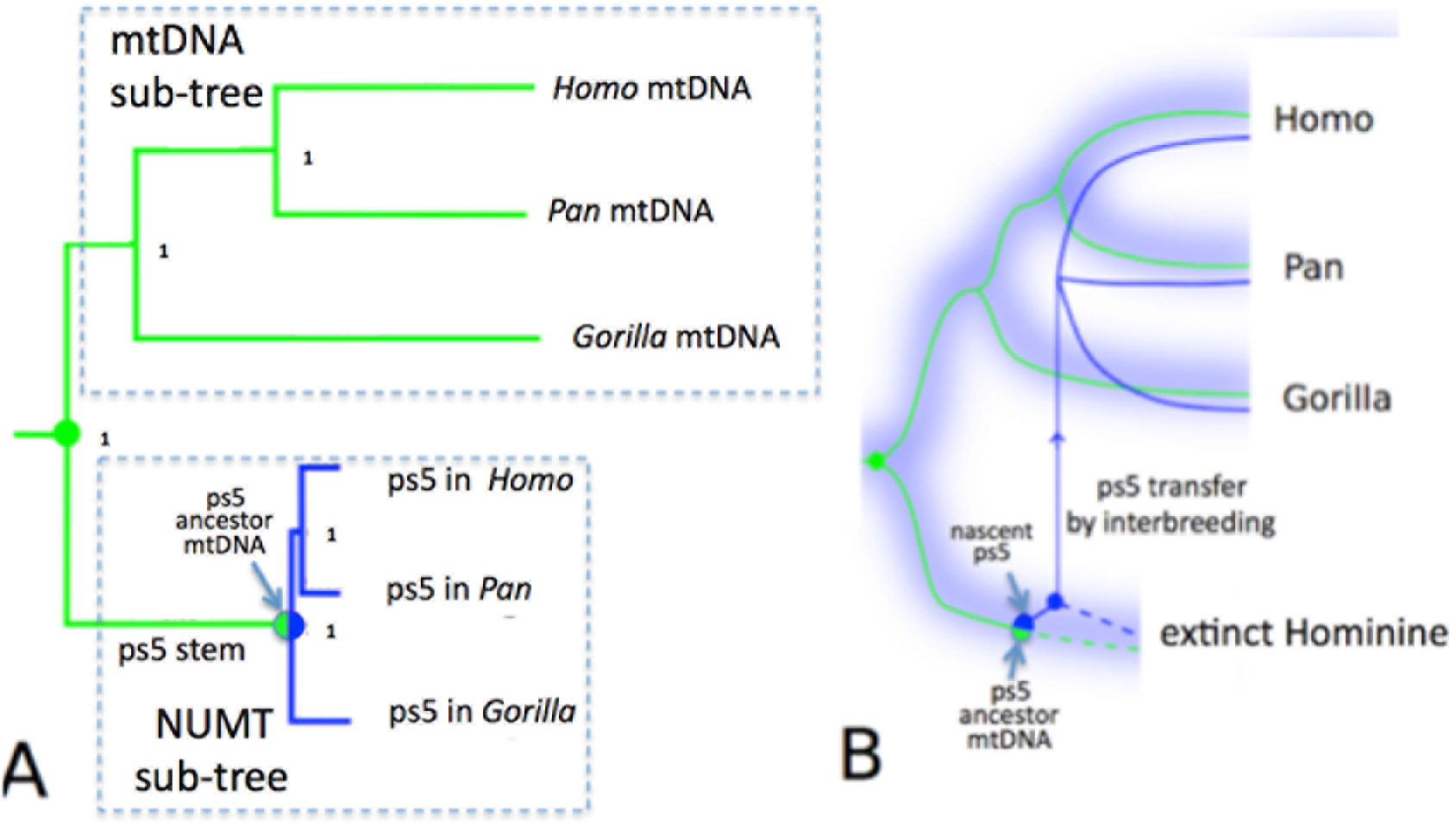
A. A joint phylogenetic tree of the hominine mtDNA and the ps5 pseudogene of mtDNA. Green and blue lines depict the mitochondrial and the pseudogene lineages respectively, diverging from their mitochondrial common ancestor (green circle). The common pseudogene stem (“Ps5 stem”) is colored green because, remarkably, mutations of the “ps5 stem” are mostly synonymous changes that must have occurred in a functional mitochondrial genome. This contrasts with a low fraction of synonymous changes in the pseudogene branches (blue). Note that the pseudogene branches are short, because of the low mutation rates in the nuclear DNA compared to mtDNA). The length of the “ps5 stem” implies ~4.5 My of evolution. The intriguing question is how did ps5 get back into the Homo/Pan/Gorilla clade after its precursor had been diverging from this clade for millions of years. Orangutan, gibbon and baboon outgroups were omitted for simplicity (see **Suppl. Note 2 and 4** for tree building approach and stability analysis). **B. Interpretation of the mtDNA/ps5 tree of Fig. 1A.** Blue hazy branches represent species, rather than individual loci, and blue haze schematically symbolizes the superposition of the phylogenetic trajectories of all the nuclear genetic loci. Because the ps5 stem branch is essentially mitochondrial, it must have been evolving within a continuous maternal lineage, which, to accommodate the very long ps5 stem, should have been diverging for ~4.5My. The long separation period implies that this maternal lineage was a part of a separate species. That species should have eventually gone extinct, and is thus labeled “extinct Hominine”. The ps5 pseudogene was created in the extinct Hominine (half blue/half green circle) and transferred to the Homo/Pan/Gorilla clade via interspecies hybridization (thin blue arrow). Solid lines represent lineages currently extant lineages, dotted lines - extinct lineages.

### The ps5 NUMT should have been transferred from a separate species

A qualitative visual comparison shows that the length of the ps5 stem is comparable in length to the *Homo* and the *Pan* mtDNA branches (**Fig. 1A**). In other words, by the time ps5 was created, the mtDNA predecessor of the ps5 pseudogene should have diverged from the *Homo-Pan-Gorilla* mtDNA almost as far as human and chimpanzee mtDNA diverged from each other. This suggests that the **ps5 stem mtDNA lineage may represent a separate, now extinct, Hominine** species. Because the ps5 sequences now reside in the *Homo*-*Pan*-*Gorilla* genomes, this **extinct Hominine should have somehow transferred these sequences to the *Homo-Pan-Gorilla* clade, which implies a distant hybridization (Figure 1B).**

**Figure 2.**
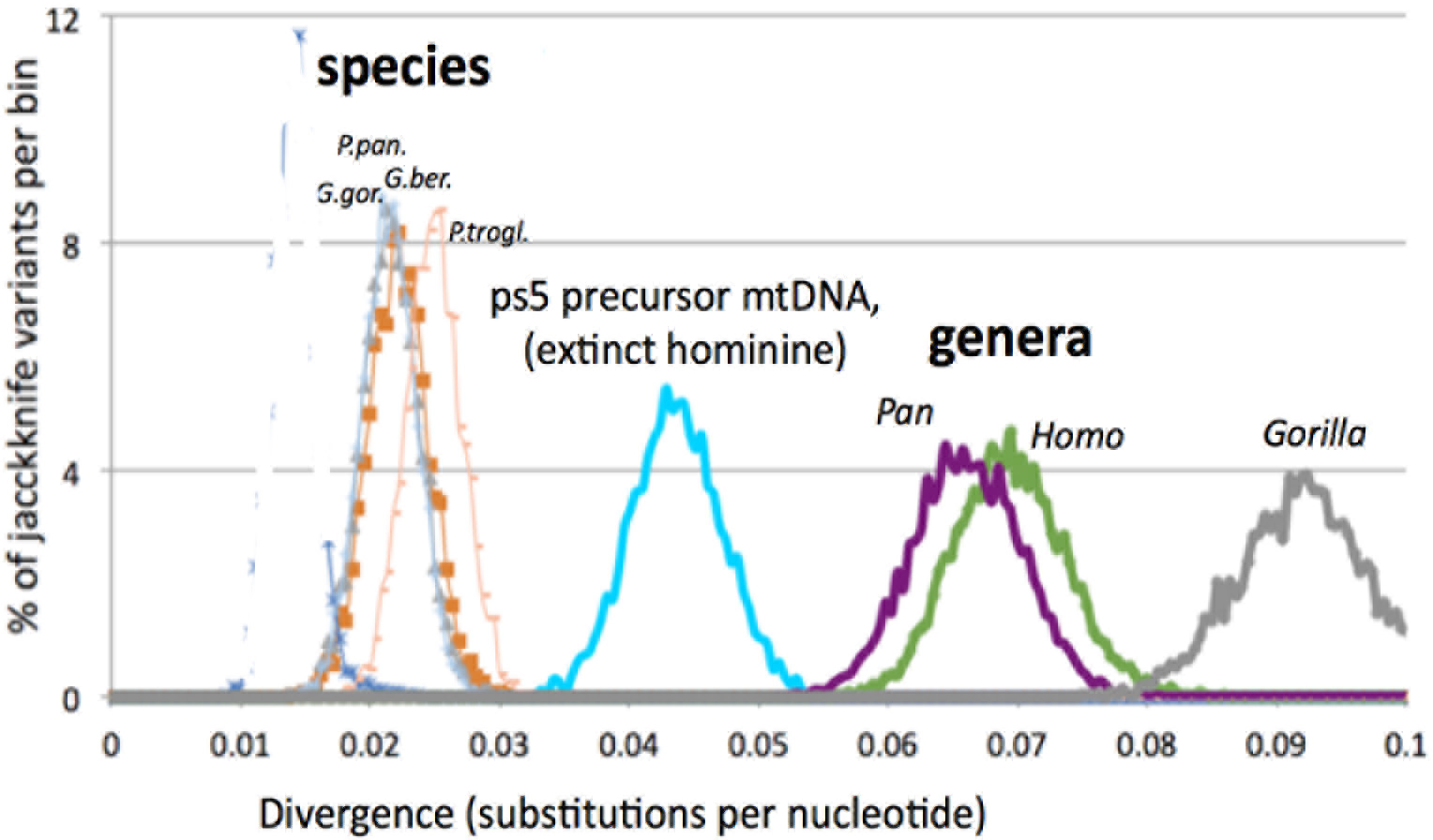
*The divergence of hominine taxa from their common ancestors with sister taxons (a.k.a. branch lengths) are compared to the divergence of the ps5 precursor mtDNA (mtDNA of the hypothetical extinct Hominine) from its common ancestor with living hominines at the time of the ps5 formation (turquoise). Note that divergence of the ps5 precursor is intermediate between the divergences of congeneric species (P.trogl. and P.paniscus; G.gor. and G.ber.) and the divergences of genera (Homo and Pan). Divergences were estimated from the common ancestor with a sister taxon (i.g. for Homo - from the common ancestor with Pan, for P.trogl. - from the common ancestor with P.paniscus, and for Ps5 precursor mtDNA - from the common ancestor with the HCG clade). The curves represent the distribution of estimates by the jackknife procedure (**Suppl. Note 3**); the ps5 data has been corrected by the fraction of mtDNA mutations in the ps5 stem (i.e. multiplied by 0.75, **Suppl. Note 6**)*

For a quantitative assessment, we estimated the mtDNA divergences within various hominine taxa and compared them to the divergence of the ps5 predecessor mtDNA using the maximum likelihood/ jackknife approach (multiple resampling of the sequence shortened by removing 50% of the base pairs at random, **Suppl. Note 3**). We used the dataset of 82 great ape mitochondrial genomes (Prado-Martinez et al., 2013), supplemented with human, Neanderthal, and Denisovan mtDNA. Importantly, this dataset was designed to represent the great ape diversity, as has been confirmed by the analysis of nuclear DNA of the same samples. Here we use the term **“% divergence”** to describe the divergence as inferred by the ML/Jackknife procedure. This measure is highly correlated with the widely accepted divergence *times* of the ape species (orangutan, gorilla, chimpanzee) and gorilla and chimpanzee subspecies-**Suppl.Note 3** and **Fig S2 therein**). The resulting distribution of jackknife estimates (**Figure 2**) shows that the ps5 stem branch (turquoise) diverged by 4.5%±0.8% from its common ancestor with *Homo-Pan-Gorilla* mtDNA by the time the pseudogene had been formed.

How much is a **4.5%** mtDNA divergence from the *taxonomic* point of view? As seen in **Figure 2**, the estimates of mtDNA divergences between congeneric species (i.e. species belonging to the same genus, represented by thin-lined curves on the left) are very well separated from the divergences between genera (thick lined purple, green, and grey curves on the right). The divergence of the Ps5 predecessor mtDNA is intermediate between the divergences of congeneric species and the divergences of genera. We thus conclude that the Ps5 precursor mtDNA and therefore its host, the hypothetical extinct Hominine, belonged to a separate species, which was significantly diverged from the human/chimpanzee /gorilla clade. Of note, divergence between the great ape mtDNA sequences increased essentially linearly with the separation time between species at about 1% per 1Myr (**Fig S2**). Thus the extinct Hominine should have diverged by about 4.5+/−0.8Myr. 4.5My is a time typically considered sufficient for significant isolation of the diverging species and such hybridization would seem impossible. However, reassuringly, a similar scenario including formation of a NUMT and its transfer to a divergent species by distant hybridization has been very recently described for Colobine monkeys (Wang et al., 2015) (see the **Discussion** section).

### Divergence of the ps5 precursor mtDNA cannot be explained by the larger size of the ancestral population

A potential alternative explanation of the high divergence of the mtDNA precursor of the ps5 pseudogene could be a high effective population size of mtDNA (Ne_mit_) of the ancestral population. In this case the expected inter-individual genetic heterogeneity can be so large that a highly genetically divergent individual could have been merely a regular member of the population, rather than an intruder from a distant species. Thus, a potential limitation of our analysis is that we used present day hominine populations as reference to assess the divergence in an ancient population, whose effective size is generally believed to be larger than that of modern great ape populations. Therefore, we asked whether a larger effective size of the ancestral population (**N*e***) rather than the taxonomic distance of the ps5 carrier could have accounted for the surprisingly high apparent divergence of the ps5 precursor mtDNA. Of note, we need to distinguish between the nuclear DNA **N*e***, ***Ne*_*nuc*_** and the mtDNA **N*e***, ***Ne*_*mit*_**. ***Ne*_*nuc*_** and ***Ne*_*mit*_** can be very different. Theoretically, ***Ne*_*nuc*_** is expected to be 4 times larger than ***Ne*_*mit*_**, but in reality the ratio depends on the particular population dynamics.

The ***Ne*_*nuc*_** of the great ape ancestral populations has been estimated recently (Prado-Martinez et al., 2013). Although the *mitochondrial* **Ne_mit_**, of the ancestral population is not known, we can use modern effective population sizes of mtDNA and nuclear DNA in order to estimate their ratio (Ne_nuc_/Ne_mit_) and, assuming that this ratio is fairly stable across the evolutionary time, to infer the ancient Ne_mit_. Thus we estimated (**Ne_mit_**) in the ancestral population in two steps: first, we determined how the mitochondrial **Ne_mit_** relates to the nuclear ***Ne*_*nuc*_** in modern hominine populations; and, second, we extrapolated that relationship to the ancestral population, assuming a constant Ne_nuc_/Ne_mit_ ratio, and finally, used this ratio to calculate the estimated mitochondrial **Ne_mit_** of the ancestral population.

We first plotted the available data on the *maximum* mtDNA divergence within present day chimpanzee and gorilla populations. As a proxy of “populations”, we used the formally accepted subspecies, conservatively assuming that individuals within a subspecies are sufficiently interconnected to be considered a population. The resulting plot (**Suppl. Note 9, Figure S4**) revealed a very weak nonsignificant correlation between Ne_nuc_ and the intraspecies mtDNA divergence. Linear extrapolation of these data to the higher Ne_nuc_ of the ancestral population shows that the anticipated mtDNA divergence in the ancestral population should have been much lower than the divergence of the mtDNA precursor of the Ps5 pseudogene (Figure S4), in accord with the “distant hybridization” hypothesis.

A possibility remains that the Ne_nuc_/Ne_mit_ was not constant and mtDNA divergence of the ancestral population was higher relative to Ne_nuc_ than that of modern populations. This would imply, however, that the ancestral population was structurally or otherwise significantly “different” from modern hominine populations in a way related to mtDNA or gender (**Supplementary Note 11**). For example, the male/female behavioral/migration patterns could have been different. Excessive divergence of mtDNA could potentially be explained by the relative immobility of females. These alternative possibilities, however, are perhaps even more peculiar and exciting than the hybridization scenario.

### Interspecies hybridization was not a unique event: evidence from NUMTs ps11 and ps7

#### Ps11: Gorilla

Ps5 is not the only mtDNA pseudogene that implies an interspecies hybridization event. Another pseudogene with similar evolutionary history has been found on Chromosome 11. Overall joint tree topology of the ps11 NUMT with mtDNA (**Figure 3**) is similar to that of ps5 (i.e. long common pseudogene stem consisting of highly synonymous, “mitochondrial” mutations and subsequent divergence among human, chimpanzee and gorilla. However, this pseudogene shows a consistently higher similarity to mtDNA of the gorilla than to that of other hominines (note a common stem segment with gorilla mtDNA in Figure 3). Note that in this case the shape of the mtDNA sub-tree poorly reflects the evolutionary history of human, chimpanzee and gorilla. This is because NUMT ps11 is homologous to the rRNA section of the mitochondrial genome. This genome segment is known to poorly reflect evolution history, possible because of excessive selection pressures.

It is tempting to speculate that the carrier of the Ps11 precursor mtDNA first belonged to the gorilla clade, then diverged into a separate lineage where it was inserted into the nuclear genome as a pseudogene, which then was transferred back to gorilla as well as to the human/chimp clade by hybridization.

**Figure 3.**
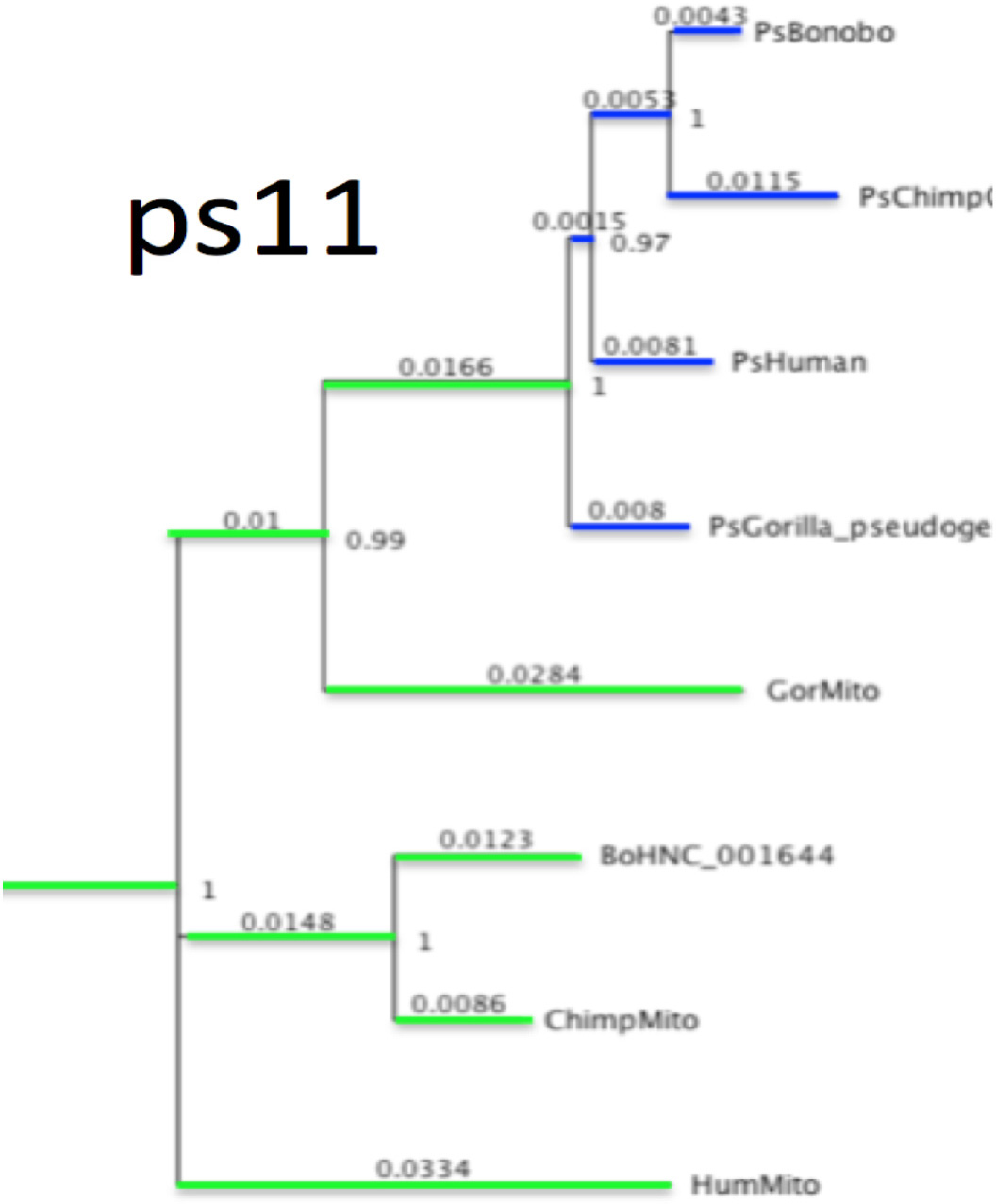
*Phylogenetics of the pseudogene Ps11 and Hominine mtDNA. PhyML GTR. Note that this tree is still under construction; there are some unresolved problems in topology. In fact, this particular region of mtDNA is generally not apt for genealogical analysis because of high conservation and high selective pressures (it is an rRNA coding region).*

Interestingly, a similar evolutionary scenario has been proposed based on the relationship between human and gorilla lice (Light and Reed, 2009). In that study, a gorilla-specific louse strain was shown to have been transferred to humans from gorillas 3.2 (+/−1.7) million years ago. Transfer of lice presumably required close persistent physical contact between members of the gorilla and our ancestors. We cannot determine with certainty the time of the ps11 transfer to the human/chimp clade because ps11 is relatively short and does not afford estimates as precise as those for ps5, but it likely falls within the anticipated broad time range of the lice transfer. It is tempting to speculate that this contact resulted in the transfer, in addition to lice, of some genes (via inter-species hybridization), and that pseudogene Ps11 is a relic of this (or similar) transfer. It is important to appreciate, however, that Ps11 mtDNA most likely has considerably diverged from gorilla by the time of becoming a pseudogene.

There is at least one more NUMT with a similar evolutionary history, which is located on chromosome 7 (**Figure 4**). This “ps7” NUMT is very old and has diverged from our lineage around the time of the Old World monkeys/apes separation. The corresponding “ps7 precursor” mtDNA has accumulated almost 9% nucleotide changes prior to its insertion into the nuclear genome. Using the same arguments as for the ps5 pseudogene we conclude that, at a certain point, there was hybridization between species with mtDNA diverged by about 9% from their common ancestor. This is a very large divergence by modern ape standards, similar to the divergence between orangutans and hominines (**Figure 4**). We will return to the plausibility of such hybridization in the Discussion.

**Figure 4.**
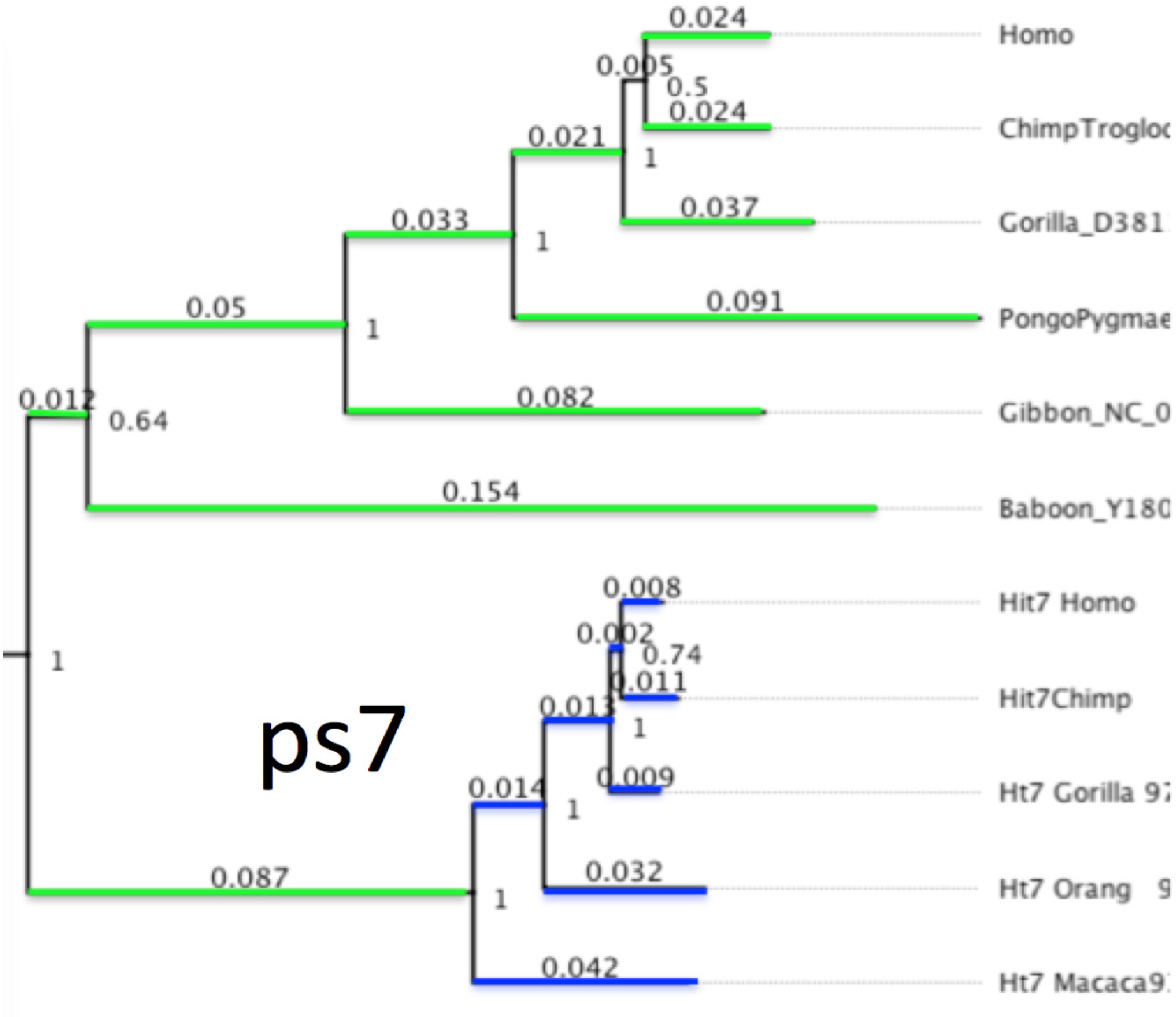
*Phylogenetics of the pseudogene Ps7 and old world monkey mtDNA.*

## Discussion

### Are such distant hybridizations plausible? Comparison to other primates

The NUMT data strongly suggests that our direct ancestors were repeatedly involved in hybridization with distant species separated for about 4.5My (ps5) and perhaps even more (ps7) prior to the hybridization event. Are such distant hybridizations at all possible? It is thought that within mammals, it takes around 2-4My, on average, to establish reproductive isolation through hybrid inviability (Fitzpatrick 2004, Evolution). However, there is considerable variation across taxa. Natural hybridization has been estimated to occur between 7 and 10 percent of primate taxa (Corteés-Ortiz et al 2007 Genetics). The majority of evidence for primate hybridization is genomic, though some phenotypic studies have also been undertaken (see discussion in Arnold and Meyer 2006, Arnold 2009, Ackermann 2010). Although the bulk of hybrids are formed between congeneric species, more distantly related intergeneric primate hybrids do occur. For example, fertile intergeneric hybrids have been documented in crosses between baboon and gelada lineages, separated for ca. 4Myr (Jolly et al., 1997). Hybridization in captivity has also occurred between rhesus macaques and baboons (so called rheboons), which diverged considerably further back in time, but they are not fertile (Moore et al 1999 AJPA). There is also evidence that some living primate species are the products of hybridization (Chakraborty et al., 2007 MPE; Osterholz et al., 2008 BMC Ev Bio; Tosi et al., 2000 MPE; Burrell et al., 2009). In one case, the kipunji (*Rungwecebus kipunji*), a baboon-like monkey, appears to be the product of hybridization ca. 600Ka between taxa whose mtDNA lineages diverged 4-6Ma, indicating that gene transfer occurred between 5.5Myr and 3.5Myr after separation of the lineages (Burrell et al., 2009). Very recently, evidence have appeared for hybridization between Asian Colobine Genera Trachypithecus and Semnopithecus, separated by the time of hybridization by about 5.5 My or more (Wang et al., 2015). Intriguingly, in this case, the evidence of the hybridization is based on a NUMT present in Semnopithecus, which is closely related to the Trachypithecus mtDNA. This implies a scenario almost identical to what we had proposed for ps5 and other pseudogenes. Given this evidence, a separation time of about 4.5Myr between the parties of the “Ps5 pseudogene transferring hybridization”, while very large, appears not unprecedented among primate lineages.

The hybridization implied by the ps7 data (9% divergence) appears too distant. It should be noted, however, that the ancestral population where this interbreeding would have taken place thrived about 30 million years ago and that little is known about its size and structure. Of note, most extant Old World monkeys practice male exogamy; if this were true for the ape/Old World monkey ancestral population, then this could have promoted high mtDNA divergence in a subpopulation, whose nDNA would not be so drastically diverged as it would be in a contemporary great ape population with the same mtDNA divergence, and thus still allowed for successful hybridization. An extreme example of such a situation is provided by the naked mole rat, where mtDNA divergence even *within* a species with rather closely interrelated nDNA reaches as high as about 5%, presumably because of extreme immobility of female queens in this eusocial rodent. Immobility of females results in an increased divergence of mtDNA, because, in this situation, local mtDNA types are rarely replaced by types from distant areas of the same population and thus can accumulate more mutations. Even with these explanatory assumptions, the divergence of ps7 pseudogene precursor is truly extraordinary. With these mtDNA pseudogenes as a lead, it would be interesting to look for other possible records (perhaps among nuclear loci) of distant interbreeding between the ancestors of our species.

### Timing of hybridization and the Fossil evidence

Of particular interest is the **time** when the asserted hybridization event took place. The phylogeny of the ps5 sequences consistently places the pseudogene insertion around the time of the *Homo/Pan* split. i.e., about 6 million years ago (**Supplemental Note 10**). In other words, the formation of the pseudogene and possibly the interspecies hybridization event took place within the Miocene epoch, when the ape lineages were diverging from each other and the human lineage was diverging from the chimpanzee clade. Intriguingly, at the terminal part of the Miocene and the early Pliocene, certain hominin fossils have been interpreted alternately as more human-like or more ape-like in different respects. For example, there is considerable disagreement about the placement of *Sahelanthropus tchadensis* (7-6Ma) on human versus ape lineages (Brunet et al., 2002; Zollikofer et al., 2006; Wolpoff et al., 2006), in part because it combines a chimp/bonobo-like cranial base and vault with more hominin-like traits, such as an anteriorly placed foramen magnum. Similarly, *Orrorin tugenensis* (~6Ma) appears to have bipedal features of the femora (Senut et al., 2001; Richmond and Jungers, 2008 Science; but see Ohman et al., 2005 Science), linking it to hominins, but ape-like dental morphology. The hominin species *Ardipithecus ramidus* (4.5-4.3Ma) possesses ape-like hands and feet, dental traits comparable to *Pan*, and cranio-dental features and bipedal capabilities that appear to link this taxon with hominins (White et al., 1994 Nature, 1995 Nature, 2009 Science). *Ar. kadabba* similarly shows a mixture of ape-like and hominin-like morphology (Haile-Selassie 2001 Nature, Haile-Selassie et al 2004 AJPA). The mosaic nature of these taxa makes them uneasy members of our clade. One possible interpretation of their taxonomic position is to place them in a separate clade of apes that shares convergent features (homoplasies) with the hominin clade (see discussion in Wood 2010 PNAS). However, it is also possible that some of these fossil specimens display mixed morphology as a result of genetic exchange between the ape and hominin lineages. This would point to a complex process of lineage divergence and hybridization early in the evolution of our lineage, with the Ps5 pseudogene representing a genomic record of such a hybridization event.

Our estimate for the timing of the alleged ps5-related hybridization event based on mtDNA/NUMTs analysis coincides with that obtained by Patterson and colleagues using nuclear DNA data i.e., “later than 6.3 Myr ago” (Patterson et al., 2008). We note however, that these two methods do not necessarily detect the same events. As discussed below, interbreeding events might have been relatively common in the evolution of our species. This coincidence may therefore indicate that such events occurred more frequently at this critical time in our evolution, during early divergence of the chimp/hominin lineages. Indeed, our preliminary data indicate that formation of mtDNA pseudogenes appear to be punctuated, potentially correlated with the epochs of speciation in the hominin lineage (Gunbin 2016). Also, while our NUMT-based analysis documents gene exchange between genetically diverse individuals with fair certainty, it provides little information on the volume of the gene flow associated with the event. In principle, almost no genes other than the ps5 itself might have been transferred from the putative extinct hominine to the HCG lineage. Therefore it is possible that the interbreeding event recorded by the ps5 pseudogene might have gone undetected by the approach used by Patterson et al., while the event they describe might have not left any NUMT record detectable by us.

### Multiple pseudogenization/hybridization events: potential positive selection of the pseudogene?

It appears that the insertion of a pseudogene coincident or swiftly followed by the hybridization of distant lineages was not a unique event in human evolutionary history. Although the consequences of hybridization vary widely, they can include the evolution of novel genotypes and phenotypes, and even new species (Arnold 1992; Seehausen 2004; Mallet 2008; Seehausen et al 2014). In the case of the human lineage, the adaptive fixation of introgressed genes appears to have occurred repeatedly, resulting in novel gene amalgamations that provided fitness advantages. For example, Neanderthal genes related to keratin production have been retained in populations living today (e.g. Sankararaman et al 2014; Vernot and Akey 2014). Similarly, genes associated with immunity (e.g. Abi-Rached et al 2011, Dannemann et al 2015) and adaptations to high-altitude environments (Huerta-Sanchez et al 2014) in living people were acquired through ancient introgression. It is therefore possible that the introgressed pseudogenes described here were linked to other genes that were themselves adaptively beneficial.

It is tempting to speculate on the possible mechanisms whereby these NUMTs got fixed in the population. Notably, the fixation process should have been rather efficient, since these pseudogenes appear to have been fixed in more than one population For example, Ps5 was independently fixed in the gorilla and the human/chimp nascent populations, which by that time were probably substantially separated. This implies that the spread of the pseudogene within and across populations might have been driven by positive selection. Interestingly, indeed, both Ps5 and Ps11 are located close to 3’ regions of functional genes (Ps7 is yet to be studied in this respect.). Insertion of an mtDNA pseudogene into the immediate vicinity of a functional gene is expected to be a strongly nonneutral mutation. In addition to a significant spatial disruption of the genome (e.g., Ps5 was a 10Kb+ insertion), the inserted mtDNA has very unusual properties, e.g., unprecedented strand asymmetry and potential for secondary structure formation (multiple RNA genes). Such an insertion is expected either to significantly alter the gene or its expression. Thus pseudogene insertion should be either highly disruptive or, rarely, significantly beneficial. The mtDNA pseudogenes that remain after millions of years may represent those rare, significantly beneficial events.

Most intriguingly, Ps11 is located within just a few hundred base pairs from 3’ transcription termination site of the RNF141/ZNF230 gene, which is essential for spermatogenesis and fertility (Zhang et al., 2001), (Song et al., 2008). It is worth noting that differences in the expression patterns of RNF141 gene were proposed to contribute to fast speciation of East Africa cichlids (PMID: 25186727), especially in the context of their strong sexual selection. Thus it is tempting to speculate that the insertion of the Ps11 pseudogene served as an expression modifier for RNF141, which resulted in increased fertility and reproductive selective advantage and eventually allowed the pseudogene to spread over the human, chimpanzee, and gorilla ancestral populations.

Interestingly, RNF141 appears to be among a few genes demonstrating a selectively driven expression shift in testis of the ancestor of hominines (PMID: 22012392). This phenomenon is perfectly in line with our hypothesis of adaptive fixation of pseudogene-induced changes in expression level of the gene. It is important to emphasize a unique nature of this expression shift – except the testis in the ancestor of hominines RNF141 did not demonstrate any adaptive expression changes in seven other investigated tissues and all other branches of the mammalian phylogenentic tree (PMID: 22012392). This strongly supports the possibility that the gene went through a phase of pseudogene-induced intense selection during the speciation of hominines. {Ackermann: 2006kc}.

### Alternative possibility: mtDNA introgression

It should be noted that a similar NUMT genealogies would have been generated in a ‘symmetrical’ hybridization scenario, where a distant species donates it’s mtDNA rather than nuclear DNA loci including the NUMT. I this scenario, NUMT is created in the acceptor species from its ‘old’ mtDNA, which later gets replaced from the population by the ‘new’ mtDNA of the introgressing species. The schematic of this event is shown in **Fig. 5A**, where it is assumed that introgression from a divergent species affects Human and Chimpanzee only (HC-introgression). The latter condition is important because if introgression affected Gorilla, the resulting shape of the mtDNA HCG tree, i.e. approximately equal H, C, and G branches) would have been incompatible with the observed (HC)G topology.

**Figure 5.**
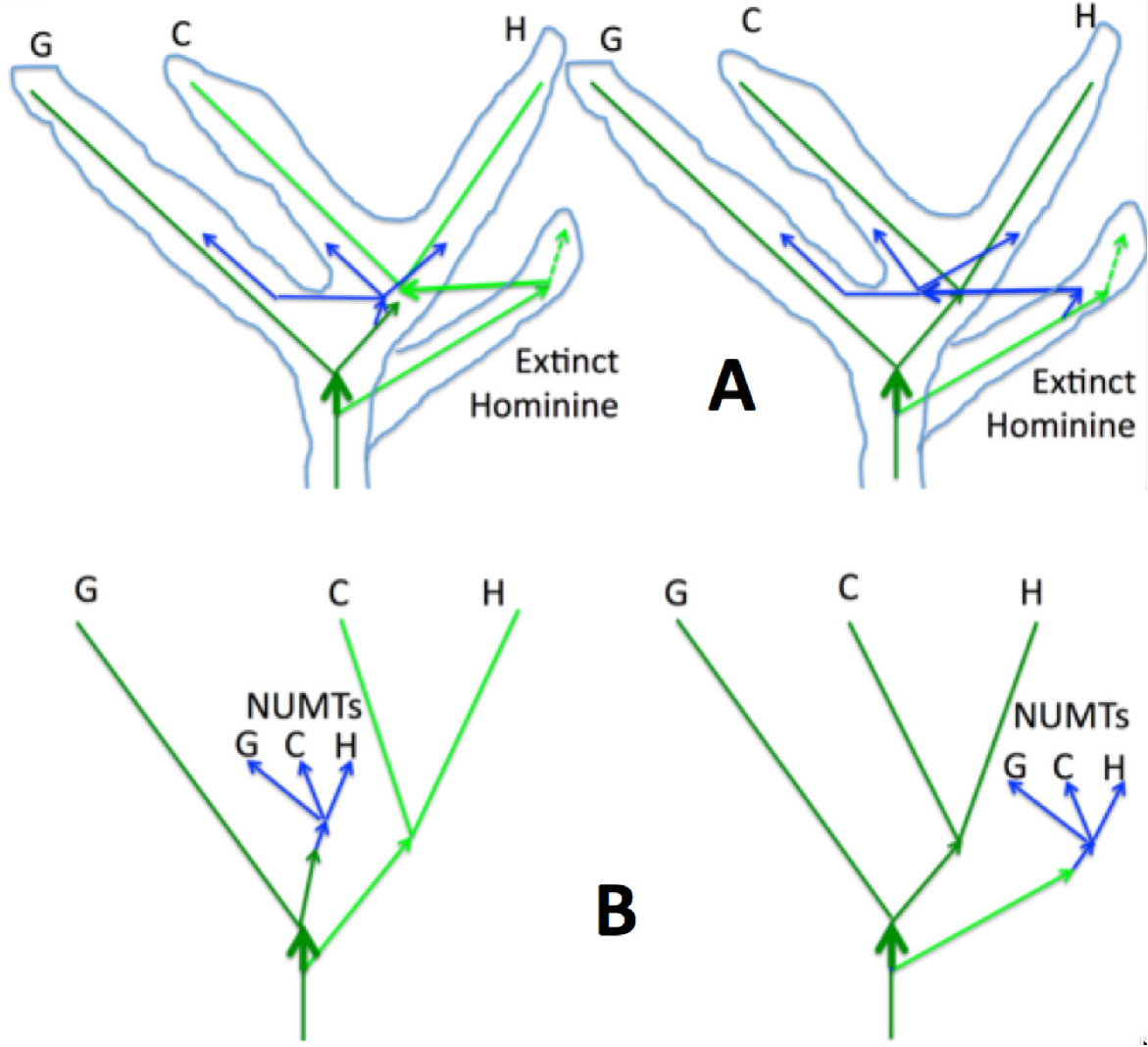
*Comparison of the NUMT transfer (right) and the mtDNA introgression (left) scenarios of hybridization. Panel **A** represents the inferred events, Panel **B** represents the resulting observed phylogenies. Blue arrows represent the NUMTs and light blue lines depict the superposition of the phylogenetic trajectories of all the nuclear genetic loci, as in Fig. 1B. mtDNA genealogies are green. Horizontal arrows represent introgression of mtDNA (green) or transfer of the NUMT locus between extinct hominine and the HCG mitochondrial (blue). Extinct hominine mtDNA lineage is neon green, the HCG mtDNA lineage is forest green*.

As shown in the figure, either of the two scenarios results in a sub-tree of NUMTs with a long stem comprised of mitochondrial mutations, as is observed in reality. The two scenarios are not fully symmetric, however. For example, the proposed mtDNA HC-introgression scenario is expected to result in a common stem between Gorilla branch and the NUMT sub-tree (thick green arrows in **Fig. 5B**). The length of this common Gorilla/NUMT stem depends on how much time elapsed between the split of the extinct hominine and the insertion of the NUMT into the nuclear genome. Intriguingly, we indeed detect various degrees of affinity to the Gorilla branch of both the ps11 and ps5 NUMTs. The significance of this preliminary observation, however, awaits further investigation. all NUMTs originating in the time interval from the split with the hypothetical extinct hominine till introgression should originate from the ‘old’ mtDNA.

## Acknowledgements

This work was funded in part by Senior Scholar award to KK from the Ellison Medical Foundation. The authors wish to thank Lukas Kuderna (Comparative Genomics Group, Institut de Biologia Evolutiva (CSIC-UPF), Universitat Pompeu Fabra and Zemin Ning (Sanger Institute, UK) for providing latest sequences of gorilla and chimpanzee prior to publication.

